# High-throughput phenomics of global ant biodiversity

**DOI:** 10.1101/2025.11.29.689474

**Authors:** Julian Katzke, Francisco Hita Garcia, Philipp D. Lösel, Fumika Azuma, Tomáš Faragó, Lazzat Aibekova, Alexandre Casadei-Ferreira, Shubham Gautam, Adrian Richter, Evropi Toulkeridou, Sabine Bremer, Elias Hamann, Jenny Hein, Janes Odar, Chandan Sarkar, Marcus Zuber, Jacobus J. Boomsma, Rodrigo M. Feitosa, Lukas Schrader, Guojie Zhang, Sándor Csősz, Minsoo Dong, Olivia Evangelista, Georg Fischer, Brian L. Fisher, Jaime Florez Fernandez, The GAGA Consortium, Fede García, Kiko Gómez, Donato Grasso, Stephane de Greef, Benoit Guénard, Peter G. Hawkes, Robert A. Johnson, Roberto A. Keller, Rasmus S. Larsen, Timothy A. Linksvayer, Cong Liu, Arthur Matte, Masako Ogasawara, Hao Ran, Juanita Rodriguez, Enrico Schifani, Ted R. Schultz, Jonathan Z. Shik, Jeffrey Sosa-Calvo, Chao Tong, Leonardo Tozetto, Seonwoo Yoon, Masashi Yoshimura, Jie Zhao, Tilo Baumbach, Evan P. Economo, Thomas van de Kamp

## Abstract

The big data era in biology is underway, but the study of organismal form has been slow to capitalize on advances in imaging and computation. Modern imaging can digitize whole organisms, but low throughput has limited the effort to document morphological diversity. Within the open science initiative ‘Antscan’, we applied high-throughput synchrotron X-ray microtomography to capture phenotypes across a diverse and ecologically dominant insect group — ants. We provide 2193 whole-body 3D ant datasets from 792 species to broadly cover the ant phylogeny with a global scope, also pairing phenomic data with genome sequencing projects.

Scans acquired with standardized parameters facilitate automated analysis and free access to data can broaden the audience and incentivize methods development. Antscan presents a scalable approach to create libraries of diverse anatomies, heralding a new era of studies on the evolution, structure, and function of organismal phenotypes.

## Introduction

The diversity of life manifests in the endless forms of organisms. Grasping this phenotypic variation is one key to understanding the evolution of organismal diversity, the interface between genomic variation and the environment, engineering principles in nature, functional traits relevant to ecosystem processes, and responses to global changes^1–3^.

Scientific collections are of fundamental value for documenting life’s diversity through space and time and form the foundation for basic and applied biodiversity science. To unlock this value, genetic sequencing, photography, and photogrammetry are established approaches for generating digital information from standing insect collections^4–7^. Truly capturing organismal form, the digitization of internal and external 3D morphological data offers the potential to expand the use of collections even further, e.g., to combine morphological and anatomical traits of form with molecular phylogenies and correlate evolutionary patterns with ecological data^8,9^. However, big data approaches to morphology and anatomy lag behind rapid advances in other digitization efforts.

X-ray computed microtomography (micro-CT) can non-destructively digitize valuable collection material in 3D^10,11^, but significant challenges must be overcome to achieve a phenomic big data revolution across the millions of animal species^9,10,12–14^. Among the major groups of animals, insects are the most abundant and species-rich class. They exhibit a vast diversity of body plans and morphological adaptations to environmental and social challenges. Insects play key functions in terrestrial ecosystems, are critical for agriculture, serve as inspiration for biomimetic design, and are important indicator organisms for measuring the effects of global change.

Towards the goal of documenting morphological diversity, synchronizing multiple facilities with conventional laboratory micro-CT scanners is one way forward, and this model has been successful for scanning vertebrates^15^. However, to image insects with conventional micro-CT, staining and/or drying is often required to discriminate between tissues^16–19^, which not only necessitates additional preprocessing steps, but alters specimens and excludes some collection material from imaging. Especially for higher resolution scans as required for smaller organisms, limits in photon flux density (the amount of radiation emitted from the X-ray source) result in long scan times, easily exceeding several hours or more. Therefore, the bottlenecks in preprocessing and throughput call for an alternative approach to achieve digitization milestones for soft and hard tissue diversity across the global insect fauna.

Further, toward the broader utilization of micro-CT data in biology, data availability represents a significant challenge since datasets are often not openly available after publication^8^ and tedious processing of 3D data is still an impediment to analyses^20^. Automation using computer vision and artificial intelligence methods can significantly reduce manual input^21,22^, which necessitates highly comparable data for training and application. Yet, in laboratory micro-CT, image properties differ significantly among specimens as staining levels and scanning conditions vary^23^, which makes applying such methods more difficult. Thus, large-scale 3D digitization efforts must also address issues of image comparability and open access.

Synchrotron-based micro-CT offers high flux density, short scanning times at high resolution, and broad contrast capabilities, particularly phase contrast, enabling visualization of soft tissue without staining. These advantages can be optimally exploited by automated high-throughput setups with robotic sample exchangers^24^. Synchrotron micro-CT has already been successful to digitize invertebrates for comparative research^24,25^ and is suitable to be extended to cover entire groups of diverse small organisms.

Within the ‘Antscan’ initiative, we generated 3D morphological data on a scale of thousands of specimens within a single week, demonstrating a powerful workflow suitable to digitize invertebrate diversity (Fig. 1). Ants occupy many ecological niches that coincide with enormous morphological and anatomical variation between and within species. They live in colonies, most incorporating worker, queen, and male castes. Many species also exhibit distinct sub-castes or more continuous variation among workers. Systematics based on robust molecular phylogenies^26–33^ and extensive ecological and behavioral research^34,35^ will enable contextualizing morphology in ecological and evolutionary research. As a globally distributed, morphologically variable and ecologically dominant group of insects, ants form an excellent basis for a pilot initiative in digitizing insect biodiversity in 3D.

**Fig. 1:**
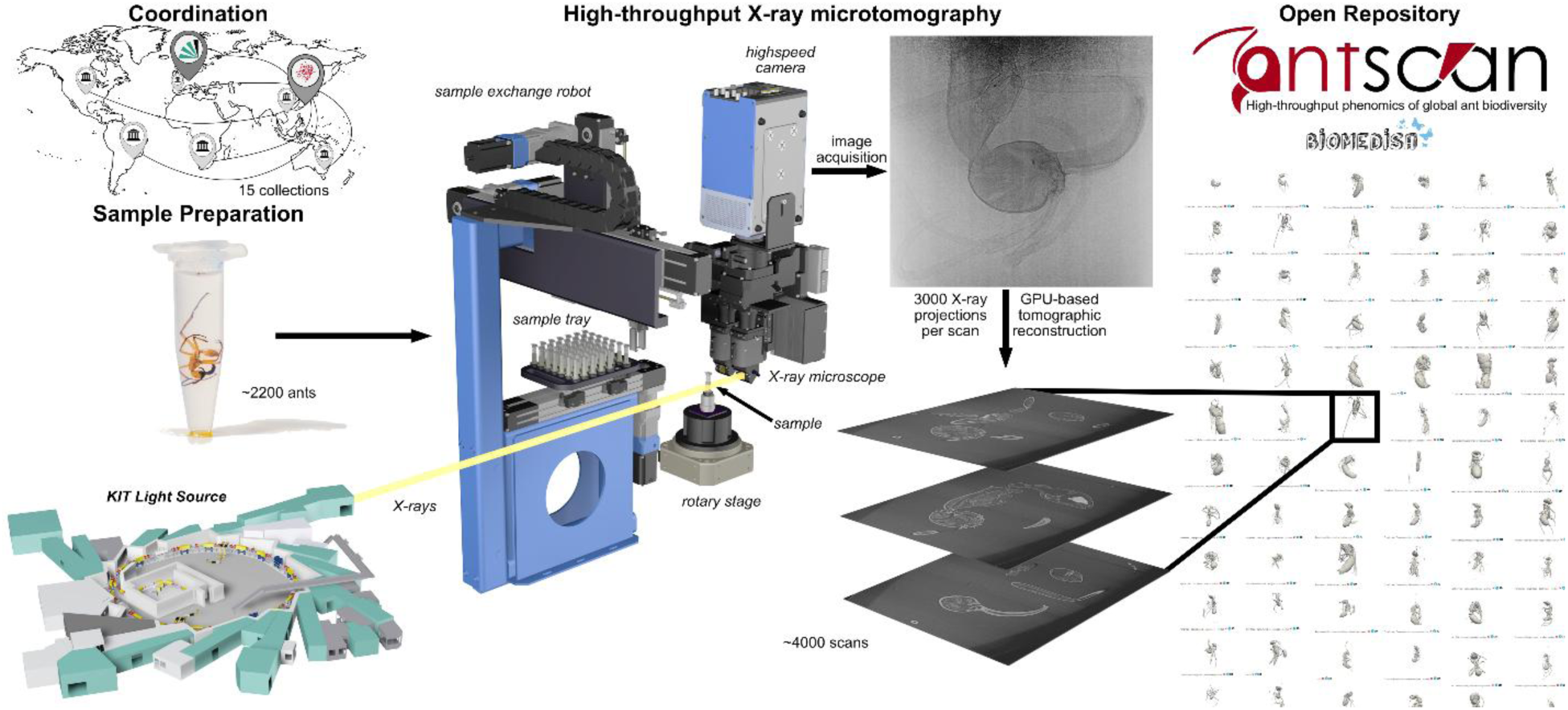
Overview of the Antscan workflow. Left Panel: Specimens were centralized and prepared before high-throughput synchrotron micro-CT scanning at KIT Light Source. Middle Panel: High-throughput synchrotron micro-CT employs the synchrotron beam as X-ray source, a sample exchange robot, a rotary stage, a high-resolution detector system, and a high-speed camera. Image stacks are then computed from the projections using GPU-based tomographic reconstruction. Right Panel: The scans are made publicly available in the online database for download, signified here by a snapshot of 3D previews.

We gathered vouchered ant samples from museums and personal collections worldwide. As soft tissue preservation is an important issue, we exclusively selected ethanol-preserved specimens. The ants were scanned *in toto* (whole-body) within ethanol using a high-throughput synchrotron micro-CT setup. Standardized scanning and 3D reconstruction protocols facilitated the creation of comparable datasets and, consequently, the application of machine learning and computer vision approaches. The processed datasets provide the foundation for the Antscan repositories, accessible via https://www.antscan.info/, which offer free access to the entire 3D image data collection of the initiative. As we synchronized our efforts with large-scale genome sequencing projects^36,37^, by targeting conspecific ants or even scanning ants from the same nest series sampled for those projects, we established a vital connection between molecular and morphological data, enhancing the scientific value of both datasets. We highlight the possibility of creating complex digital 3D models by semi-automatic segmentation and show how simple screening through a vast number of datasets can quickly recover patterns in morphological and ecological adaptations.

Antscan not only serves as a vast open resource for promoting research on the morphology and anatomy of ants but also establishes a design that can be adapted for other lineages of small organisms across the tree of life. By digitizing and democratizing an important aspect of scientific collections while strictly keeping the data tied to their sources and collectors (see Supplemental Table S2), these valuable resources become accessible for analysis worldwide. Making the exceptional phenotypic diversity of ants available in 3D thus unlocks a previously inaccessible wealth of information for a broader audience, enabling comprehensive analyses and contributing to the establishment of online repositories for researchers, educators, or nature enthusiasts. Our vision is to embed this methodology into the activities of the global scientific community.

## Results

### Phylogenetic coverage

We designed specimen sampling within Antscan to be phylogenetically broad, to represent species-poor clades, to include rarely collected species, and to sample multiple representatives of highly diverse genera. Datasets cover 14 out of the 16 currently recognized extant subfamilies, and ants were identified to 212 out of about 345 presently recognized genera^30,38^ (Fig. 2). These 212 genera incorporate more than 90 % of all described ant species. Out of 2193 scanned ants, 1711 specimens were identified to 659 species, and within 482 not fully identified ant specimens, we currently recognize at least 133 distinct morphospecies, resulting in at least 792 ant species represented in our total dataset. Antscan includes 1671 workers, 291 queens, and 220 males.

**Fig. 2:**
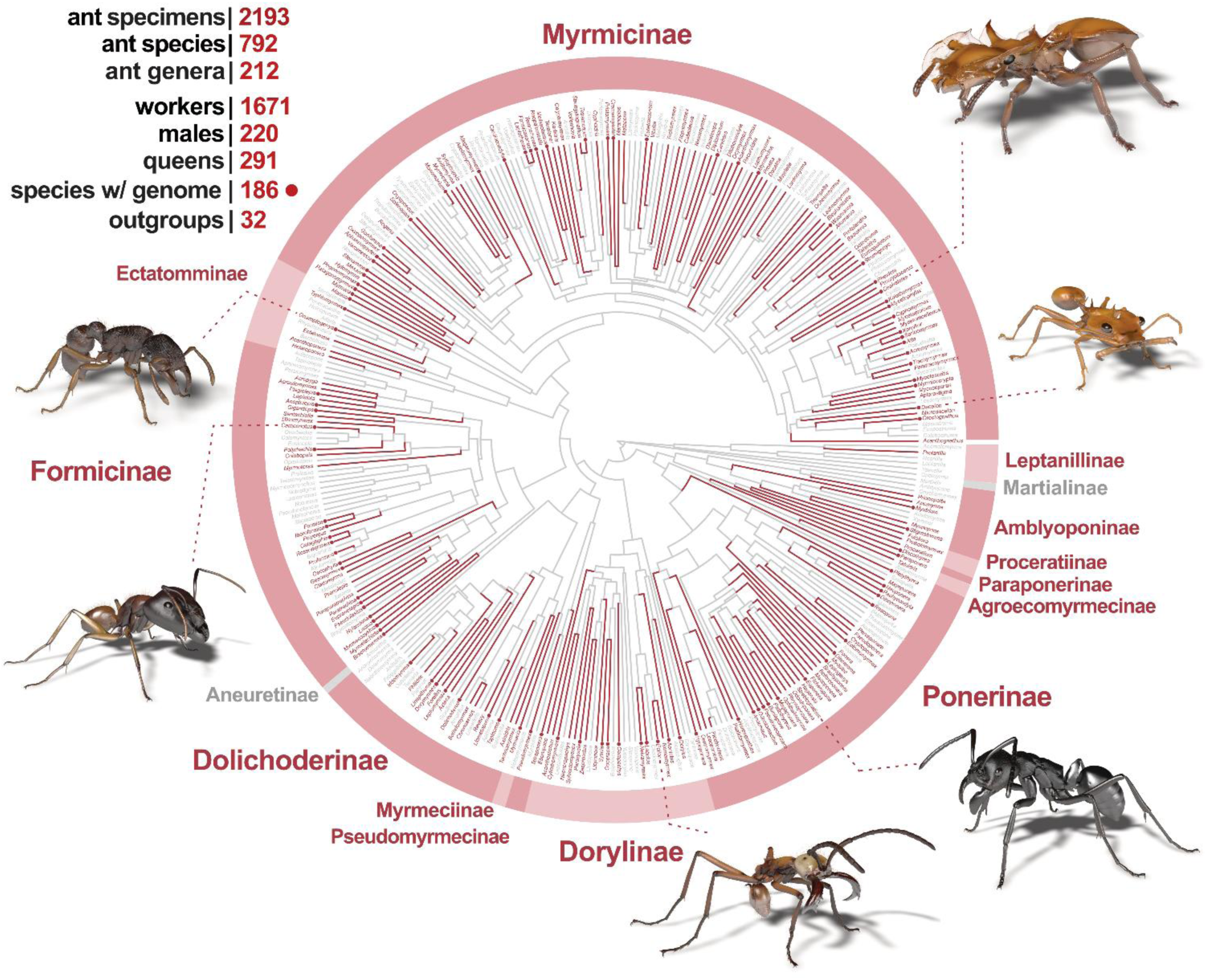
The diverse set of species and genera across the ant tree of life covererd by Antscan. The ant phylogeny with genera sampled in Antscan displayed in red. A dot at the end of a genus branch indicates that at least one genome from a species within that genus is sequenced. Examples of 3D models derived from the micro-CT scans shown clockwise from the top: *Cephalotes* sp., *Daceton armigerum*, *Streblognathus peetersi*, *Eciton hamatum*, *Camponotus* sp., *Gnamptogenys* aff. *continua*. The phylogenetic tree is based on Economo et al.^33^ with missing genera added through taxonomic affinities or other published phylogenies^26–32^.

Within the worker caste, we identified 18 ants as explicitly being media workers and 125 as major workers, with more polymorphic ants yet to be delimited. In addition to the ant datasets, we provide 32 scans of non-ant wasps for outgroup comparison. For the three most species-rich ant genera *Camponotus*, *Pheidole,* and *Strumigenys,* which incorporate more than 850 species each, we sampled 37, 33, and 39 species, respectively. Important currently monotypic genera like *Apomyrma*, *Santschiella*, or *Tatuidris* are included, variations of peculiar morphologies, such as various trap-jaw ants or multiple turtle ant species are covered, and globally important species like fire ants or Argentine ants were sampled, allowing for the investigation of a plethora of research questions building on the global diversity of ants.

In coordination with genome sequencing projects^36,37^, 585 of the scanned specimens will be associated with genomic data, of which 157 specimens are even taken from the same nest series as the sequenced specimens. On the species level, the pairing of Antscan and genomic data translates into 186 species. As many taxa have further been covered by published molecular phylogenies, with Antscan, we aim to form a platform for future comparative research on the genomic basis of phenotypic variation and diversification in invertebrates.

### Tomographic data

By employing high-throughput synchrotron micro-CT at two beamlines of the KIT Light Source, we generated *in toto* tomograms for 2193 ant plus 32 outgroup specimens. Three magnification settings with corresponding fields of view were employed to optimally cover the wide range of ant sizes in the collection (Fig. 3, Material and Methods). Specimens exceeding the field of view vertically were scanned in several height steps.

**Fig. 3:**
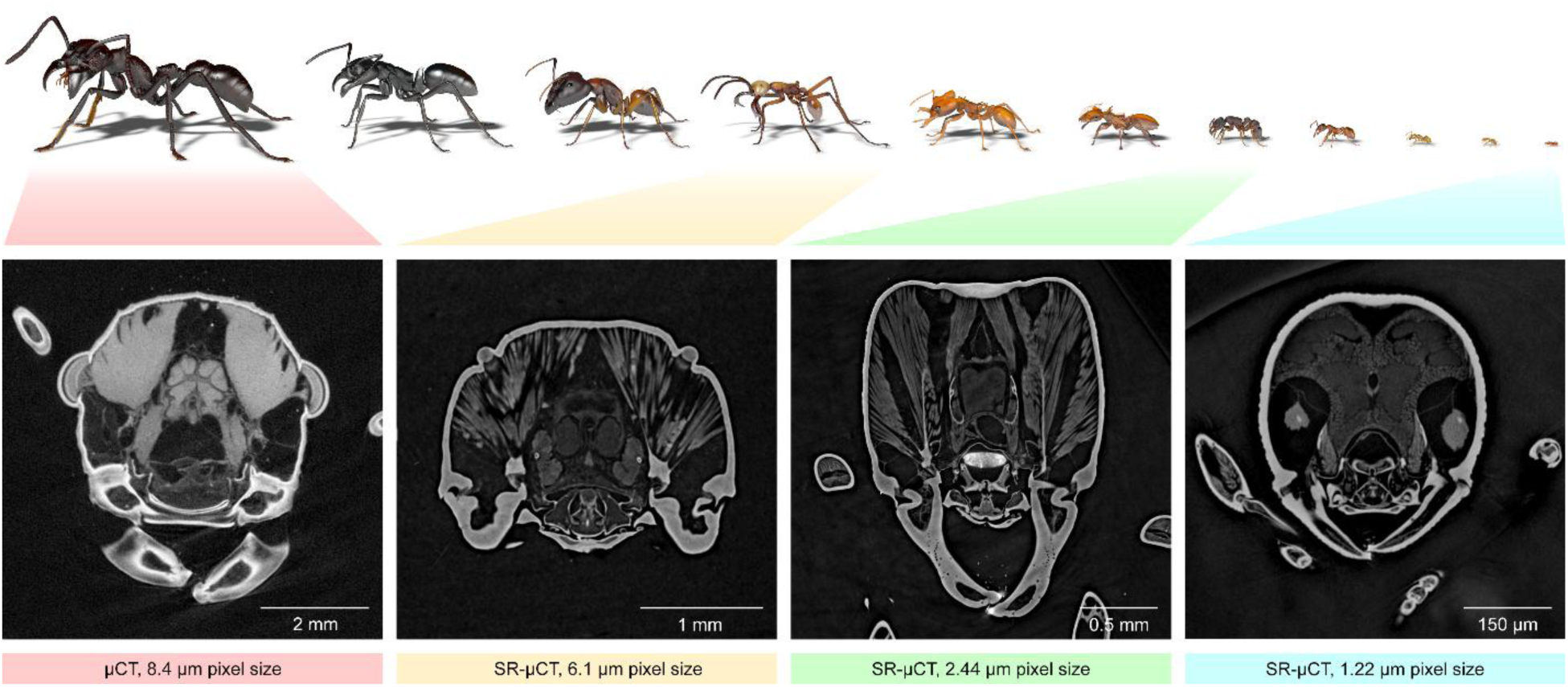
Exemplar images representing different magnifications for different-sized ant workers. Top: 3D models of different ant workers scanned within Antscan depicted to illustrate operational scales. When scaled to an A4 page, the ants appear in their original size. **Bottom:** Image slices from the heads of four ant species acquired with different magnifications. From left to right: *Paraponera clavata* stained with iodine and scanned using laboratory micro-CT as one of the largest ants that did not fit inside the largest field of view of the synchrotron setup; *Eciton hamatum* subsoldier as a large ant; *Gnamptogenys* aff. *continua* as a medium-sized ant; *Discothyrea sexarticulata* as one of the smallest ants.

In addition to absorption contrast, synchrotron measurements also exploit phase contrast due to the partial transverse coherence of the radiation. Phase contrast arises during the free-space propagation of the transmitted wave field, which already after short propagation distances enhances the tissue boundaries in the measured images (so-called edge enhancement^39^). This is particularly important for soft tissues such as muscles, which absorb almost no radiation. A phase retrieval algorithm then converts the edge enhancement into distinguishable tissue contrast. To obtain good overall contrast for both higher absorbing parts like the exoskeleton and soft tissues, the primary datasets of the Antscan database are blended volumes, composed of one volume obtained by standard 3D reconstruction of the measured images and the other by additional phase retrieval applied prior to the 3D reconstruction.

As an exception to the blended-volume datasets that comprise most of the Antscan scans, we deviated for 132 previously iodine-treated ants and the six largest ants we gathered. Due to the strong absorption of their soft tissues, most of the iodine-stained specimens were reconstructed without phase-retrieval. For the six largest specimens, which exceeded even the largest field of view available at the synchrotron setup in diameter, we performed iodine-staining and scanned them with a laboratory micro-tomograph. Although differing in imaging, we could incorporate these specimens in all downstream processing steps.

While the original tomograms are saved as 32-bit data, we converted and further processed the tomograms as 8-bit TIFF stacks. When required, we used a computer-vision workflow to merge individual height steps automatically into a combined volumetric dataset, showing the entire ant body. Employing a neural network using Biomedisa^21,40^, we further crudely segmented datasets automatically and cropped all tomograms by removing background and thus reducing file size considerably. This automated processing of all data underlines the suitability of high-throughput synchrotron micro-CT for future large-scale analyses using computer vision and machine learning.

Since the data acquisition and processing parameters were identical for the sample subsets scanned at a given magnification at a beamline (see Methods), the reconstructed gray values of the corresponding tissues within these subsets are equivalent (except for few iodine-treated samples), ensuring comparability and facilitating the application of machine learning and computer vision approaches.

For quality assurance, we visually inspected all scans to check for specimen preservation quality. Most ants were well preserved, but unavoidable when drawing from standing collections, but some showed severe damage from soft tissue shrinkage and decay, probably caused by DNA extraction, exposure to air during transport, handling, or storage. However, as the rigid exoskeleton was generally not affected, many morphological or morphometric analyses can still be applied. Overall, given the preservation quality, we conclude that both short- and long-term stored ethanol-preserved material is generally suitable for non-destructive high-throughput synchrotron micro-CT to digitally preserve 3D anatomy of invertebrates indefinitely.

Since samples were prepared and positioned manually beforehand and scanning proceeded automatically, the partial truncation of some datasets could not be avoided. Legs and antennae of larger specimens within a magnification category were particularly prone to be outside the field of view during scanning. Future developments towards improved robotic setups and imaging pipelines that can autonomously recognize the individual position of specimens during measurements and readjust scanning positions accordingly will likely resolve such minor issues.

### The Public Antscan Repositories

All tomograms acquired within Antscan and their associated metadata are provided publicly in open repositories that can be accessed via the Antscan website (https://antscan.info) The interface for the primary, interactive repository (https://biomedisa.info/antscan) was created on the image analysis and segmentation platform Biomedisa^40^. A search function offers the possibility to explore the database for specific keywords in the metadata. Individual datasets are represented on the top page by 3D surface previews based on neural network automatic segmentation. The ant datasets then feature more previews for image slices and an overview of the insect habitus as an interactive 3D model of the surface mesh. Apart from the tomographic data, each ant is accompanied by comprehensive metadata, including taxonomic ranks, ecological parameters, and unique specimen identifiers (Supplemental Table S2). We also provide information on specimen locality, further geographical details, habitat, and whether a sequenced genome is associated with the specimen. Metadata further provides the information required to identify and acknowledge curators, other contributors, and host institutions responsible for the individual specimens.

The public Antscan database infrastructure is curated, aims to provide an interactive experience, and is sustainable. Integration into the Biomedisa interface offers the possibility for registered users to apply methods for semi-automatic and automatic volumetric image segmentation with the Biomedisa app, which can be directly applied to the datasets. Datasets that are being processed can be easily shared with other Biomedisa users. In addition to Biomedisa, we ensure the sustainability of the entire Antscan database by providing another long-term-secured location for all files including metadata on the RADAR4KIT repository of KIT (https://radar.kit.edu/radar/en/search?query=antscan). The chosen format of a mirrored, open database allows for smooth phylogeny/taxonomy-based online navigation and assures public long-term availability of the Antscan data.

### Exemplary use cases

To demonstrate the wealth of information contained in an individual ant scan, we segmented exoskeletal elements, muscles, nervous tissues, sting apparatus, and digestive tract of a sub-soldier of the South American army ant *Eciton hamatum* (Fig. 4). Such segmented anatomical features can, e.g., be extracted and quantified for analyses, visualized as renderings (Fig. 2, 3 & 4a), animated (see Extended Data Video E1), and 3D-printed. Detailed digital examinations and the extraction of morphological characters of single specimens already do and will likely lead to new discoveries.

**Fig. 4:**
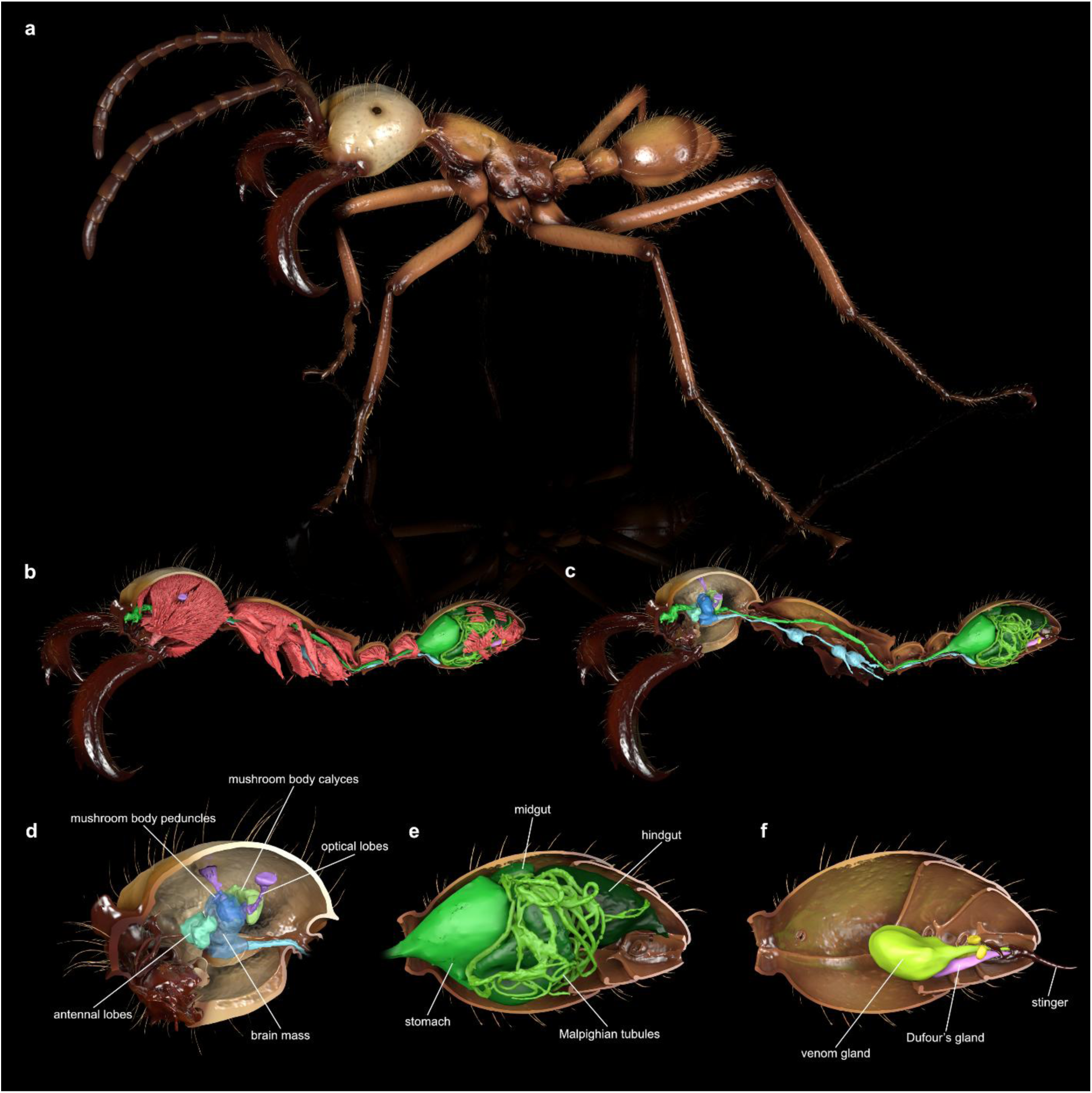
Renderings of an exemplary Antscan specimen (*Eciton hamatum* sub-soldier CASENT0744582) showing the segmented cuticle and tissues representative of the level of detail captured with synchrotron micro-CT. **a:** Full habitus of the ant with an animated, more life-like pose and colors inspired by photographs. **b:** Cuticle cut at the sagittal section revealing internal tissues with muscles in red occupying most of the internal space in an ant’s body. **c:** Removing the muscles reveals the digestive tract (green) and the nervous system (blue). **d-f:** Zoomed-in renderings focusing on the ant brain, gut, and sting apparatus, respectively.

Moreover, the vast amount of available specimen data facilitates large-scale comparative studies, e.g., to identify similarities and differences between lineages and to trace the evolution of traits throughout the ant tree of life. In this context, not all scientific cases require extensive segmentations of individual ants. Simple screening of the datasets may already quickly clarify whether morphological features are present or absent in ant species or entire lineages. Here, this is shown on the example of biomineral armor, an unusual insect trait first described for *Acromyrmex echinatior*^41^ and previously unknown in other ants. The Antscan data immediately revealed that this trait is more widely distributed. A conspicuous highly absorbing layer on top of the cuticle confirms that biomineral armor is in fact common among fungus-growing ants (Myrmicinae: Attini) and scattered across the different agricultural systems that evolved in these clades^42,43^ (Fig. 5). Within the attine ants, it appears to be absent in the genera *Atta*, *Kalathomyrmex*, and *Mycocepurus*, which is consistent with an inferred secondary loss of biomineral armor^41^. We have not found evidence of this trait in any species outside the Attini.

**Fig. 5:**
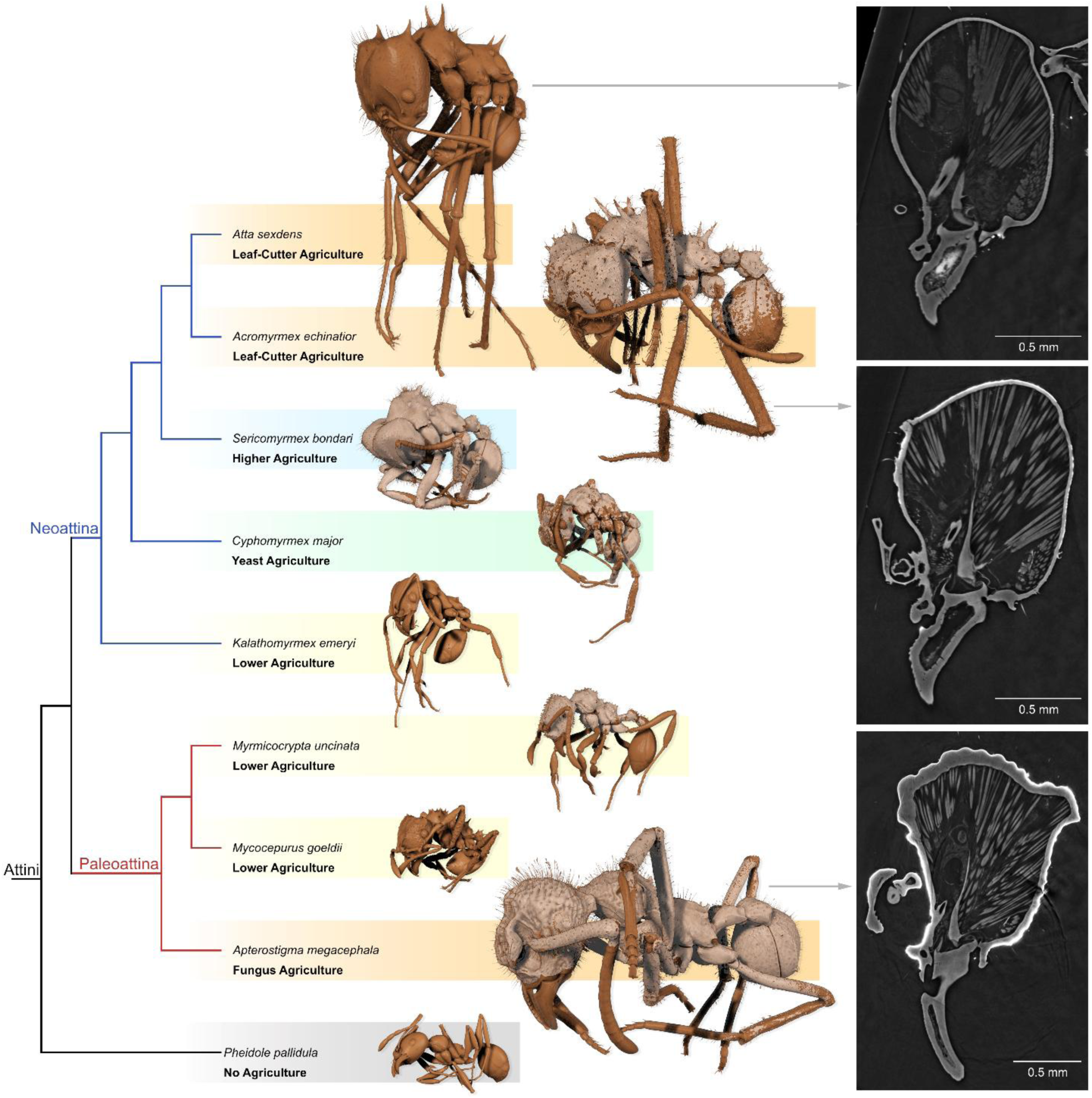
Trait recognition with Antscan on the example of biomineral armor. Comparative screening revealed that biomineral armor is very common among fungus-growing ants and was found in several attine species. The slices through the tomograms (right) show the armor as a thin but distinct coat of highly absorbing (brighter) material on top of the cuticle. In the simple thresholded 3D renderings (left) it is visualized as a beige color.

This example illustrates the potential for standardized high-throughput imaging datasets to enable testing evolutionary and ecological hypotheses at larger scales without the acquisition of new data.

## Discussion

With Antscan, we present a scalable design to meet the demands for large-scale 3D digitization of phenotypes from biodiversity collections. It hinges on first centralizing specimens to fit them to the high-throughput imaging setup, then scanning in a condensed timeframe, reconstructing and processing CT-scans to finally provide them in an open-access format. We explicitly address points of criticism in the accessibility of and credit for digitized biodiversity; our database is open, and the metadata identifies all contributors^8,11,44^.

Inherently, high-throughput synchrotron micro-CT effectively overcomes the bottleneck of acquisition times in micro-CT. It is fast and provides well-resolved morphological datasets with high contrast for both hard and soft tissues if specimens have been properly preserved. Including file transfer, which currently constitutes a technological bottleneck in our scanning setup, Antscan moved at a pace of ∼25 scans per hour. Similarly resolved scans of ants using conventional micro-CT would take about 12 hours each on average^45,46^, with the additional delay of individual scan setups. Extrapolating this to the 4,010 individual scans in Antscan, it would have taken one laboratory micro-CT scanner operated around the clock more than six years to obtain a similar dataset.

The Antscan workflow implementation does demand considerable effort and coordination. A collaborative approach among global collections stakeholders is necessary, including meticulous management of institutional and governmental requirements, such as in specimen transfer. This engagement, however, translates into scientific value through accurate documentation of specimens and accountability for contributors. The laborious process of sorting and preparing specimens to fit the robotic setup is indispensable for achieving optimal results. After raw data acquisition, the reconstruction of tomographic volumes poses a significant technical challenge, being time-consuming and computationally demanding. Nevertheless, experiences from Antscan will help drive future workflow improvements, such as the generalization and automation of tomography pipelines or the automated post-processing of imaging data. The total storage demands for the present datasets and raw data exceed 100 Terabytes, thereby necessitating long-term commitment from storage-hosting institutions. Further, it is imperative to ease data processing for comparative research, which currently represents a major bottleneck^20,47^.

Computer vision methods, particularly neural networks, have the potential to reliably identify tissues in micro-CT data, as shown recently for ant and bee brains^21,22^ and as we show here with the automated generation of whole-body segmentations. When developed further, the Antscan project design can be scaled to hundreds of thousands and potentially millions of specimens. Its imaging principles allow both in-depth examination of individual specimens and fast screening and automated analysis of large data subsets, which will be crucial in future research based on digital morphology.

High-throughput synchrotron micro-CT in close collaboration with collectors and collection managers promises a key solution for digitizing 3D anatomy across small invertebrates. A remaining significant obstacle to the broader use of synchrotron tomography is the limited beamtime availability at synchrotron facilities. Most of them offer user service based on peer-reviewed proposals, and obtaining beamtime remains competitive. Automation of synchrotron facility imaging stations through robots and the resulting high-throughput setups are also developed to very different degrees. Moreover, the homogeneously illuminated field of view at many synchrotron X-ray imaging beamlines is restricted, which impedes the scanning of larger specimens. However, current developments, such as longer beamlines^48^ or Bragg crystal optics^49^, and novel sample exchange systems point towards a broader range of organism and collection sizes to be imaged with comparable synchrotron micro-CT setups.

The scientific community benefits from a rapidly growing collection of commercial and open-access software that facilitates analysis and visualization of 3D volumetric datasets^50,51^. Volume renderings based on grey values can be employed to quickly generate impressive 3D views^52,53^, while surface meshes allow digital dissections and the creation of interactive 3D models^45,54^.

Segmentation remains the most common feature extraction method for CT scans^51^, and the contrast properties across Antscan datasets facilitate this process for users. Due to the diversity of specimens sampled, Antscan offers the possibility to visualize external and internal morphology on larger comparative scales than previously possible with tomographic data, especially for insects.

In contrast to physical specimens, digital information can be directly accessed from anywhere in the world, enabling immediate and simultaneous access by researchers, artists, and the public, thus allowing wider and more equitable engagement with biodiversity. Antscan is a scalable approach to digitize small-bodied organisms in 3D to make the “micro” world more accessible. Like a genome, a 3D scan contains deep information about an organism, but obtaining quantitative information from micro-CT scans remains challenging. We will need new bioinformatics based on automated image analysis to fully unlock the potential of databases like Antscan, but recent developments in this area show promise^21,22^. The Antscan initiative aims to empower and encourage people around the world to engage with and incorporate highly resolved ant morphology and anatomy into their science. With the approach described here and further developments in imaging technology, bioinformatics, and artificial intelligence on the horizon, it is time for the study of phenotypes to take its place alongside other big data endeavors in biology.

## Materials & methods

### Material

We gathered ants preserved in ethanol from museum-, university-, and private collections. 132 specimens were previously stained with iodine (see Supplemental Table S2). At OIST, we checked the vials, chose apparently well-preserved specimens, transferred them to fresh 99% ethanol, assigned unique specimen IDs, and, depending on their size, stored ants individually in 0.2 ml, 0.5 ml, and 1.5 ml plastic reaction tubes that would fit in the robotic setup at KIT. We extracted and databased label data from the specimens to link relevant ecological metadata to the specimens. We then preliminarily sorted specimens into scan trays by size, each tray containing 48 vials for automated scanning. To optimize the sorting step prior to scanning, specimens within one tray should have about the same size within the tube. That way, they can be assigned conveniently to the different available magnifications, and if necessary, the number of necessary height steps can be estimated for the tray. We labeled vials with a code to trace each specimen back to its respective metadata. Tube labels were then the basis for file names during scanning. To make this physical identifier as legible and permanent as possible, flat snap-cap plastic tubes with ideally a machine-written code attached to the top of the lid have proven useful. All specimens are being stored indefinitely at OIST or have been returned to their owners or managing institutions.

### High-throughput synchrotron X-ray microtomography

The specimens were scanned within two campaigns at the IPS Imaging Cluster at the KIT Light Source. Due to the available access, two different beamlines were used. During both campaigns, the same high-throughput tomography experimental station^55^ was employed, ensuring identical detector systems, available magnifications, scanning resolutions and sample exchange robotics (Advanced Design Consulting USA, Inc.). This guarantees the comparability of datasets at a given magnification setting within each measurement campaign In the first campaign, a set of samples was investigated using a parallel polychromatic X-ray beam produced by a 1.5 T bending magnet, spectrally filtered by 0.5 mm aluminum to remove low energy components from the beam^56^. The resulting spectrum peaked at about 17 keV, and a full width at half maximum bandwidth of about 10 keV. In the second campaign, the samples were scanned by using a beam diffracted by a Double Multilayer Monochromator (DMM). The measurements were performed at a magnetic field of the wiggler of 2.7 T, yielding a maximum flux density at 16 keV with an energy bandwidth of 2%. To reduce the heat load on the DMM, the beam was pre-filtered with pyrolytic graphite.

Depending on their size, ants were scanned with magnifications of 10x, 5x, or 2x, resulting in effective pixel sizes of 1.22 µm, 2.44 µm, or 6.11 µm, respectively. An air-bearing rotary stage (RT150S, LAB Motion Systems) served for sample rotation. A fast indirect detector system consisting of a scintillator (10x: 13 µm LSO: Tb; 5x: 25 µm LSO: Tb; 2x: 200 µm LuAG: Ce), a double objective white beam microscope^57^ (Optique Peter) and a 12-bit pco.dimax high-speed camera (Excelitas PCO GmbH) with 2016 × 2016 pixels was employed. For each scan, 200 dark field images, 200 flat field images, and 3000 equiangularly spaced radiographic projections in a range of 180° were taken with exposure times between 6.25 milliseconds and 25 milliseconds, resulting in scan durations between 21 and 85 seconds. If the ants were too large for the field of view, additional height steps were scanned. In total, we acquired 4010 scans (10x: 165; 5x: 3026; 2x: 807; laboratory micro-CT: 12) for 2193 individual ants plus 32 outgroup Hymenoptera (10x: 149; 5x: 1831; 2x: 239; laboratory micro-CT: 6).

The control system ‘concert’^58^ served for automated data acquisition and online reconstruction of tomographic slices for data quality assurance. Online and final data processing included dark and flat field correction and, if applicable, phase retrieval of the projections based on the transport of intensity equation^59^. X-ray beam parameters for algorithms in the data processing pipeline were computed by syris^60^, and the execution of the pipelines, including online tomographic reconstruction, was performed by the UFO framework^55^. The final 3D tomographic reconstruction was performed by tofu^61^ and additionally included ring removal, 8-bit conversion, and blending of phase and absorption 3D reconstructions. Phase reconstruction was not performed for the previously iodine-stained specimens if their soft tissues still absorbed strongly. For the automatic processing of large sample series, our reconstruction software was extended by a sample and rotation axis finding procedure, which used a contrast measure to locate the top and bottom of a sample in 2D projection. For these two positions, the program reconstructed several slices with different axis positions and computed a gradient-based measure to determine the correct axis. From having the correct axes for the top and bottom parts, the program could then detect whether the tomographic axis was perfectly aligned with the pixel columns and, if not, corrected for such bias.

Reconstructed tomogram image stacks are ordered from bottom to top. As conventions on how to read in data differ between software applications, the end-user must confirm that the orientation of the scan is correct. Otherwise, the scan may appear mirrored, and it will be necessary to flip the scan to re-establish correct orientation.

### Laboratory X-ray microtomography

The six largest specimens were scanned in the X-ray laboratory for computed tomography and laminography of the IPS Imaging Cluster. For these, we employed a microfocus X-ray tube (XWT-225, X-RAY WorX, Garbsen, Germany), producing a conical, polychromatic X-ray beam from a solid tungsten target. We acquired the radiographic projections with a flatpanel detector (XRD 1621 CN14 ES PerkinElmer, Waltham, USA) featuring 2048 x 2048 pixels with a physical pixel size of 200 µm and a DRZ+ scintillator. Using the ‘concert’ control system, all components are positioned by a custom-built manipulator system. For the scans, the X-ray tube was operated with an acceleration voltage of 60 kV and a target power of 15 W. A total of 2048 projections were acquired over an angular range of 360°. Each frame was exposed for 4 s. We placed the samples 69.7 mm away from the source and the detector 1630.5 mm downstream of the samples, resulting in a magnification of 24.4 and an effective pixel size of 8.2 µm. We performed two separate scans for each specimen to cover the whole animal. The 3D volumes were reconstructed with tofu^61^.

### Post-processing of tomographic data

We employed an additional computer vision pipeline to, if necessary, merge height stages, generate 3D stacks of reduced file size cropped to the ant, 3D surface mesh models, and sampled down image stacks for 2D preview. We first stitched all individual CT scans for a single specimen to merge height steps based on Matte’s Mutual Information metric in a multi-resolution framework to sample exhaustively at lower resolutions before applying a Gradient Descent Optimizer at finer scales implemented in Simple-ITK^62,63^. To stitch the height steps together, we first resampled the images to corresponding positions retrieved from registration and then, for merging, computed a weighted average gradient for the overlapping region where each of the two scans was given higher influence for the result the closer it was to the overlapping slice. For all datasets, we then used Biomedisa’s deep learning feature^21^ to generate automated full-body segmentations. We trained this network with annotated data consisting of 12 randomly selected scans, which we manually pre-segmented, followed by Biomedisa’s semi-automated segmentation^64^. We used the trained network to segment whole bodies of all datasets and subsequently dilated these segmentations to mask a region around the ants. We then used the minimum and maximum indices of the segmentations in all image planes to crop the volumetric images. With this workflow, we reduced the size of the online database by about 70 %. Upon completion of merging and cropping, we retrained the neural network with 34 whole-body segmentations, of which 27 were used for training and 7 for validation (Supplemental Data S3). With this new deep neural network, we performed whole-body segmentations for all datasets.

Using largest-islands selection to remove small parts, we isolated the ant to produce 3D surface meshes in STL format and rendered those in Paraview^65^ for a 2D preview of the 3D habitus of the specimen. Finally, we sampled all stacks down to obtain a preview representation of the entire image stack.

### Further post-processing of exemplary data sets

For slices in Figures 3 and 5, we opened tomograms in Fiji^66^ or Amira to extract images with adjusted contrast to maximize visual tissue differentiation and separation from background. For the surface renderings shown in Figures 2-4, we employed Amira or Slicer 3D for pre-segmentation of whole exoskeletons, individual sclerites, and selected organs. Pre-segmented labels served as input for semi-automatic segmentation with Biomedisa^40,64^. When using Biomedisa, we imported the results back into Amira and corrected minor errors. For the 3D models shown in Fig. 5, we employed Amira’s threshold tool to segment cuticle (threshold 105-255) and biomineral armor (200-255). We converted final label fields into polygon meshes, exported meshes as OBJ files, and reassembled and smoothed them in CINEMA 4D R20. We also employed CINEMA 4D for artificial coloring, final image rendering, and animation.

### Biomedisa instance of the Antscan database

The tomographic datasets of Antscan are stored and archived at KIT’s Scientific Computing Center and linked with the Biomedisa platform. The metadata are stored in Biomedisa’s MySQL database and backed up on an external storage system^67^.

As datasets will be processed to generate new data and metadata of specimens that may change in the future (such as updated taxonomic status), we designed the database to enable such changes. All datasets can be freely accessed and downloaded. Selected registered Biomedisa users may act as administrators and can thus assign repository access to other users and edit and annotate database entries. All registered users can perform semi-automated and automated segmentation analysis and may append their own data, such as segmentation results, to the specimens.

### RADAR4KIT instance of the Antscan database

In addition to Biomedisa, we ensure the sustainability of the entire Antscan database by providing another long-term-secured public location for all files including digital object identifiers (DOIs) and metadata on the RADAR4KIT repository of the Karlsruhe Institute of Technology (https://radar.kit.edu/radar/en/search?query=antscan). We uploaded post-processed tomograms from KIT’s Scientific Computing Center to RADAR4KIT using a dedicated API and included metadata. We appended DOIs to the metadata in other locations to further enhance the accountability of each dataset generated within Antscan.

## Author Contributions

J.K. and F.H.G. contributed equally to the work. E.P.E., and T.v.d.K. jointly supervised the work.

J.K., F.H.G., T.F., P.D.L., J.J.B., R.M.F., L.S., G.Z., T.B., E.P.E., and T.v.d.K. conceived and conceptualized Antscan.

F.H.G., A.C.-F., J.J.B., S.C., M.D., O.E., R.M.F., G.F., B.L.F., J.F.F., The GAGA Consortium, F.G., K.G., S.d.G., D.G., B.G., P.G.H., R.A.J., R.A.K., R.S.L., T.A.L., C.L., M.O., H.R., J.R., E.S., L.S., T.R.S., J.Z.S., J.S-C., C.T., L.T., S.Y., M.Y., G.Z., J.Z., and E.P.E. contributed material and contributed to the specimens included in Antscan. J.K., F.H.G., F.A., A.R., L.A., S.G., E.T., and T.v.d.K. prepared material for the imaging experiment.

J.K., T.F., A.R., E.H., J.O., M.Z., and T.v.d.K. performed the micro-CT experiments.

J.K., T.F., P.D.L., S.B., E.H., J.H., A.M., C.S., M.Z., and T.v.d.K. participated in reconstructing and/or processing the data.

J.K., T.F., P.D.L., and C.S. built online database infrastructure.

J.K., P.D.L., A.R., A.C.-F., L.A., S.G., and T.v.d.K. performed 3D reconstructions to include in the manuscript.

J.K., E.P.E., and T.v.d.K wrote the initial manuscript and created the figures. All authors extensively commented on and revised the manuscript at all stages.

## Data availability

All processed Antscan tomograms are stored at the Large Scale Data Facility (LSDF) of KIT’s Scientific Computing Center (SCC) and integrated into Biomedisa for public access (https://biomedisa.info/antscan/). Additionally, all datasets are archived at the RADAR4KIT repository (https://radar.kit.edu/radar/en/search?query=antscan). Both versions of the database and all metadata and DOIs linked to datasets are accessible via https://antscan.info. All metadata is also presented in the supplemental information to this publication. Raw X-ray projections and all other image files are further stored at KIT and will be made accessible upon reasonable request.

## Code availability

We provide code and instructions to post-process micro-CT data on Github (https://github.com/julesforfools/Antscan). We further provide the deep neural network used to segment, crop and visualize ant datasets directly as a supplementary file (Supplemental data S3) in H5 format. This trained network is intended to be reused or retrained within the Biomedisa deep learning framework (https://github.com/biomedisa/).

## Acknowledgements & Funding

Research at KIT was supported by the projects HIGH-LIFE (05K2019) and SMART-Morph (05K2022) via the German Ministry for Research and Education (BMBF). We gratefully acknowledge the data storage service SDS@hd supported by the Ministry of Science, Research and the Arts Baden-Württemberg (MWK) and the German Research Foundation (DFG) through grant INST 35/1503-1 FUGG. J.K., F.H.G., F.A., A.R., A.C.-F., L.A., S.G., E.T., A.M. and E.P.E were supported by funding from the Okinawa Institute of Science and Technology Graduate University. To F.H.G., we acknowledge the Japan Society for the Promotion of Science (JSPS) grants-in-aid KAKENHI grants no. 18K14768 and 21K06326. P.D.L. has received funding from the Australian Research Council via the ARC Training Centre for Multiscale 3D Imaging, Modelling, and Manufacturing (M3D Innovation, project IC 180100008). To L.A, we acknowledge the Japan Society for the Promotion of Science (JSPS) grants-in-aid KAKENHI grants No. 22KJ3077. S.C. was supported by HUN-REN Hungarian Research Network and by the National Research, Development, and innovation Fund Grant No. K 147781. R.M.F. was supported by the Conselho Nacional de Desenvolvimento Científico e Tecnológico (CNPq) (grant 301495/2019-0). L.S. was funded by the Deutsche Forschungsgemeinschaft (DFG, German Research Foundation) – 502787686. P.G.H was funded in part by the Critical Ecosystem Partnership Fund (CEPF). The Critical Ecosystem Partnership Fund is a joint initiative of l’Agence Française de Développement, Conservation International, the European Union, the Global Environment Facility, the Government of Japan, and the World Bank. A fundamental goal is to ensure civil society is engaged in biodiversity conservation. B.L.F. acknowledges support through NSF Grant DEB-1932467. D.A.G. has benefited from the equipment and framework of the COMP-HUB and COMP-R Initiatives, funded by the ‘Departments of Excellence’ program of the Italian Ministry for University and Research (MIUR, 2018-2022 and MUR, 2023-2027). B.G. acknowledged funding from the Environment and Conservation Fund (Hong Kong), project Nb. ECF 137/2020. R.A.K. was supported by FCT (UIDB/00329/2020; DOI 10.54499/UIDB/00329/2020). T.A.L. acknowledges support through NSF Grant NSF IOS-2128304. T.R.S. and J.S.C. were supported by U.S. National Science Foundation grant DEB 1927161.

We greatly appreciate Christian Peeters and Philip S. Ward for their guidance during the planning of the Antscan initiative. We want to thank Angelica Cecilia for her great assistance at the IMAGE beamline and for contributing text to the methods section. The original 3D models of the equipment shown in Fig. 1 were kindly provided by Ursula Herberger (KIT-IBPT; KIT Light Source), AVS|US Inc. (sample exchange robot), LAB Motion Systems (rotary stage), Optique Peter (detector system) & Excelitas PCO GmbH (high-speed camera). We acknowledge the KIT Light Source for provision of instruments at their beamlines and we would like to thank the Institute for Beam Physics and Technology (IBPT) for the operation of the storage ring, the Karlsruhe Research Accelerator (KARA).

